# Adaptive mechanoproperties characterize glioblastoma fitness for invasion

**DOI:** 10.1101/2020.06.17.156406

**Authors:** Pascale Monzo, Michele Crestani, Yuk Kien Chong, Katharina Hennig, Andrea Ghisleni, Qingsen Li, Cristina Richichi, Paolo Maiuri, Martial Balland, Michael P. Sheetz, Giuliana Pelicci, Beng Ti Ang, Carol Tang, Nils C. Gauthier

**Affiliations:** IFOM - the Firc Institute of Molecular Oncology; Via Adamello, 16, 20139 Milan, Italy; Neuro-Oncology Research Laboratory, Department of Research, National Neuroscience Institute, Singapore; Université Grenoble Alpes, F-38042 Grenoble, France; CNRS UMR 5588 LIPhy, F-38041 Grenoble, France; Department of Experimental Oncology, IEO, European Institute of Oncology IRCCS, 20139 Milan, Italy; Department of Biochemistry and Molecular Biology, University of Texas Medical Branch, Galveston, TX, USA; Mechanobiology Institute, National University of Singapore, Singapore 117411, Singapore; Department of Translational Medicine, Piemonte Orientale University “Amedeo Avogadro,” Novara, Italy; Singapore Institute for Clinical Sciences, Agency for Science, Technology and Research (A*STAR), Singapore; Department of Neurosurgery, National Neuroscience Institute, Singapore; Duke-National University of Singapore Medical School, Singapore; Department of Physiology, Yong Loo Lin School of Medicine, National University of Singapore; Division of Cellular and Molecular Research, Humphrey Oei Institute of Cancer Research, National Cancer Centre, Singapore; School of Biological Sciences, Nanyang Technological University

## Abstract

Glioblastoma are heterogeneous tumors composed of highly invasive and highly proliferative clones. Heterogeneity in invasiveness could emerge from discrete biophysical properties linked to specific molecular expression. We identified clones of patient-derived glioma propagating cells that were either highly proliferative or highly invasive and compared their cellular architecture, migratory and biophysical properties. We discovered that invasiveness was linked to cellular fitness. The most invasive cells were stiffer, developed higher mechanical forces on the substrate and moved stochastically. The mechanochemical-induced expression of the formin FMN1 conferred invasive strength that was confirmed in patient samples. Moreover, FMN1 ectopic expression in less invasive clones increased fitness parameters. Mechanistically, FMN1 acts from the microtubule lattice, counteracting microtubule bundling and promoting a robust cell-body cohesion leading to highly invasive hurdling motility.

**One Sentence Summary:** FMN1 increases cell mechanics and invasiveness.

---

Glioblastoma (GBM) is the most common and the most aggressive of all primary brain tumors, with a high recurrence rate and a median survival of 14 months (*1, 2*). To this day, no cure for glioblastoma is available. Various cells, all from brain origin, can give rise to glioblastoma, including neural stem cells, multipotent neural progenitor cells, oligodendroglial precursor cells, astrocytes and neurons (*1*). These cells are able to invade and destroy most regions of the brain, leading to an increasing debilitation due to neurodegeneration (*1*). Treatment includes surgery, radiotherapy and chemotherapy (*1, 2*). Despite these aggressive treatments, glioblastoma tumors always relapse, leading to the rapid death of the patients. This is mostly due to the fact that glioblastoma cells are highly invasive, spreading rapidly in various areas of the brain, and escape treatment (*1, 2*). To this day, no treatment has been found that efficiently blocks glioma invasion, probably because these cells display specific motility modes that need to be precisely defined in order to identify valid molecular targets. Unlike many metastatic cells, glioma cells do not use the blood circulation to disseminate. Instead, they migrate actively onto specific paths such as the white matter tracts and the surface of the brain blood vessels, also known as “secondary structures of Scherer” (*3–9*). Several studies have shown that microfabricated linear tracks, including nanofibers, linear grooves, microchannels, and linear micropatterns can mimic brain linear tracks and potentiate glioma motility (*2, 10*), especially if laminin is used (*11–14*).

Few targets have been involved in the glioma linear migration process. They include cytoskeletal molecules such as actin, microtubules, myosin II, the formin FHOD3 as well as STAT3 signaling molecules (*13, 15*). However, most of these studies have been performed using *in vitro* glioma models grown with serum that have lost the specificity of human glioblastoma invasive cell populations. Cell lines isolated from patient samples and grown as tumor-spheres without serum constitute today’s best biological tool to study glioblastoma cell biology. These cells are able to grow and form a GBM tumor when injected in mouse brain (and hence are called <u>h</u>uman <u>g</u>lioma <u>p</u>ropagating <u>c</u>ells, hGPCs) (*16, 17*). Depending on the cell line, the tumor will grow and spread at different speeds and with different patterns, ultimately killing the animal at different times. These hGPC are mostly analyzed in bulk, as tumorspheres. Their invasive abilities are studied *in vivo* as xenografts, with little distinction between cells that invade actively or exhibit expansive growth. Knowing the molecular mechanisms that distinguish active invasion versus expansive growth could bring considerable advance to our knowledge of glioblastoma pathology and could help designing new tools to tackle this disease.

We compared the motility behavior of 3 hGPCs (namely NNI-11, NNI-21 and NNI-24) isolated from patients and carefully chosen, out of a vast library of xenografts, for their different invasive and proliferative behaviors. In xenografts, the NNI-11 developed tumors that were circumscribed (i.e., expansive growth), and fast-growing (Fig. 1A and fig. S1). In contrast, the 2 other cell lines, NNI-21 and NNI-24 were diffusive. The main differences between the NNI-21 and NNI-24 tumors were the fact that NNI-21 were fast-growing and led to the death of the animals faster than the NNI-24, which grew slower (Fig. 1A-C and fig. S1).

**Fig. 1.**
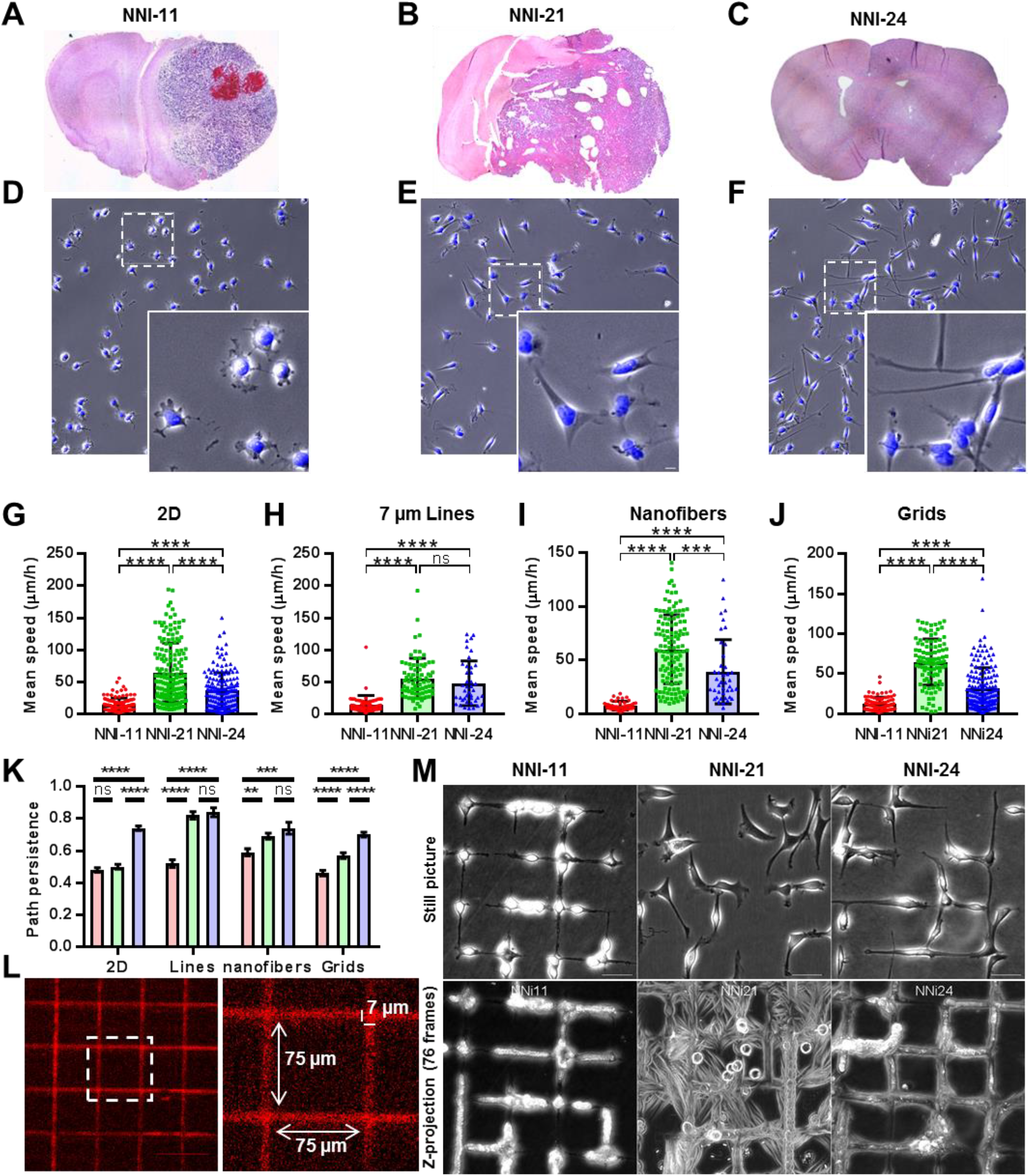
Hurdlers are able of rapid directional changes evidenced by gridded micropatterns. (**A-C**) Xenografted mouse brain slices. Mice were injected in the brain with hGPCs and sacrificed after 3 months (NNI-11), 2 months (NNI-21) and 4.5 months (NNI-24). (**D-F**) Images and zooms of cells on 2D laminin-coated surface. Cells are stained with Dapi (scale bar is 10 μm). (**G-J**) Mean speed (μm/h) of the 3 hGPCs over 6 hours on laminin-coated substrates: (**G**) 2D (n= 119, 167, 168); (**H**) micropatterned lines (7 μm width) (n=61, 77, 41); (**I**) nanofibers (1.3 μm diameter)(n=46, 127, 41); (**J**) grids (7 μm width, 75 μm gap) (n=87, 102, 168). (**K**) Average of path persistence over 6 hours on the 4 settings (red is NNI-11, Green is NNI-21, blue is NNI-24). (**L**) Image, zoom and specifications of the gridded micropatterns used throughout the study. (**M**) Up: Typical snapshot of hGPC migrating on grids; down: projection of 76 frames corresponding to 7.6 h of a movie of cells migrating on grids. n = number of cells, for speed measurement and path persistence in each setting, is given for NNi11, NNI-21 and NNI-24, respectively. Error bars are S.E.M.

## Mimicking migration complexity unveils peculiar hurdling motility in the most aggressive clone

We analyzed the migration of these hGPCs using substrates of increasing complexity, all coated with laminin, since this matrix protein is highly present around brain blood vessels and promotes glioma motility (*13, 18*). Moreover, our 3 hGPC showed better adhesiveness and motility on laminin than on other tested substrates (Movie S1). On glass-bottom dishes, 3 different migratory behaviors and cell shapes were detected (Fig. 1D-G, K, fig. S2 and movie S2). The NNI-11 were small, displayed short and thin protrusions, and were poorly motile, in accord with their non-diffusive behavior, while the diffusive NNI-21 and NNI-24 were actively migrating. The NNI-21 were fast (mean speed average = 60 μm/h) and poorly persistent. They displayed a large cell body and short processes. In contrast, the NNI-24 were slow (mean speed average = 30 μm/h) and highly persistent (Fig. 1D-G, K, fig. S2 and movie S2). They had a small cell body and one single long leading process (Fig. 1D-G, K, fig. S2 and movie S2). However, these 2D approaches poorly represent the brain 3D environment. Since, in the brain, glioma cells mainly follow linear tracks, we used different types of laminin-coated linear tracks to analyze their migration. On micropatterned lines, both NNI-21 and NNI-24 were able to migrate great distances at similar rates with persistence (Fig. 1H, K, fig. S2 and movie S2). On suspended nanofibers that mimic 3D linear tracks (*19*), similar observations were made: both hGPCs displayed good motility and persistence (Fig. 1I, K, fig. S2 and movie S2). When encountering crossed fibers, the NNI-21 readily changed direction, often hanging on several fibers at the time, but without limiting their migration ability. The NNI-24, in contrast, remained steady on a single fiber and rarely switched from one fiber to another. To better exemplify this phenomenon and quantify it, we printed crisscrossing strips of 7 μm in width that formed X intersections that could accentuate the ability of these cells to change direction while keeping a linear motility (Fig. 1J-M, fig. S2 and movies S2-S3). On these grids, the difference between the NNI-21 and NNI-24 cell lines was remarkable, especially when the whole temporal stacks were projected, as displayed in Fig. 1M. In this case, NNI-21 fully displayed their stochastic, jumpy motion, cutting angles when meeting crossroads, while the NNI-24 slowed down and followed a single track over time (movies S2 and S3). Because of these phenotypes, NNI-21 were named ‘hurdlers’ and NNI-24 were name ‘gliders’. The NNI-11 were non-motile on the 4 systems.

## Hurdlers are stiffer and display numerous dynamic adhesions

We hypothesized that these differences in motility behavior were related to the mechanical properties of the cells and measured several mechanical parameters. Adhesion topology and dynamics were measured by TIRFM (<u>T</u>otal <u>I</u>nternal <u>R</u>eflection <u>F</u>luorescence <u>M</u>icroscopy) and FRAP (<u>F</u>luorescence <u>R</u>ecovery <u>A</u>fter <u>P</u>hoto-bleaching) in cells stained for endogenous vinculin (Fig. 2A-J) or overexpressing mcherry-vinculin (Fig. 2N, fig. S3 and movie S4). A cross-correlation program was developed to track the adhesion dynamics and calculate their half-life (see Materials and Methods). Traction force parameters were extracted from cells seeded on laminin lines printed on 5kPa acrylamide gels, as described in Hennig (*20*) (Fig. 2K-M). Young’s modulus was extracted using AFM (<u>a</u>tomic <u>f</u>orce <u>m</u>icroscopy), from cells seeded on 2D or grids (Fig. 2O-P). We found that several cell mechanical properties correlated to their motility behavior. Traction forces of the 2 motile hGPCs, hurdlers and gliders, were higher than 10 nN at the front and the back of the cell (Fig. 2K-M) and clearly much higher than those of the non-motile NNI-11. The adhesion topology and dynamics could explain why the hurdlers were prone to change direction more frequently than the 2 other cell lines. In these cells, when seeded on grids, adhesions were distributed as large patches simultaneously onto several lines (Fig. 2B) and were highly dynamic (adhesion half-life was 1.44 min with a confidence interval ranging from 1.007 to 2.07 min)(Fig. S3 and movie S4) with a high rate of turnover of the vinculin in the adhesions (Fig. 2N and fig. S3A-B). In comparison, gliders displayed fewer adhesions, which were less dynamic (adhesion half-life was 3.155 min with a confidence interval ranging from 2.466 to 4.166 min) and the non-motile NNI-11 displayed extremely small and poorly dynamic adhesions (adhesion half-life was longer than the movie length of 20 min)(Fig. 2A-J,N, fig. S3 and movie S4). In addition, the hurdlers were the stiffest (Fig. 2O-P) compared to the 2 other cell lines, with a Young’s modulus of 500 Pa (versus 400 and 300 for the NNI-24 and NNI-11 respectively) indicating that a greater cytoskeletal cohesion was linked to a better ability to change shapes.

**Fig. 2.**
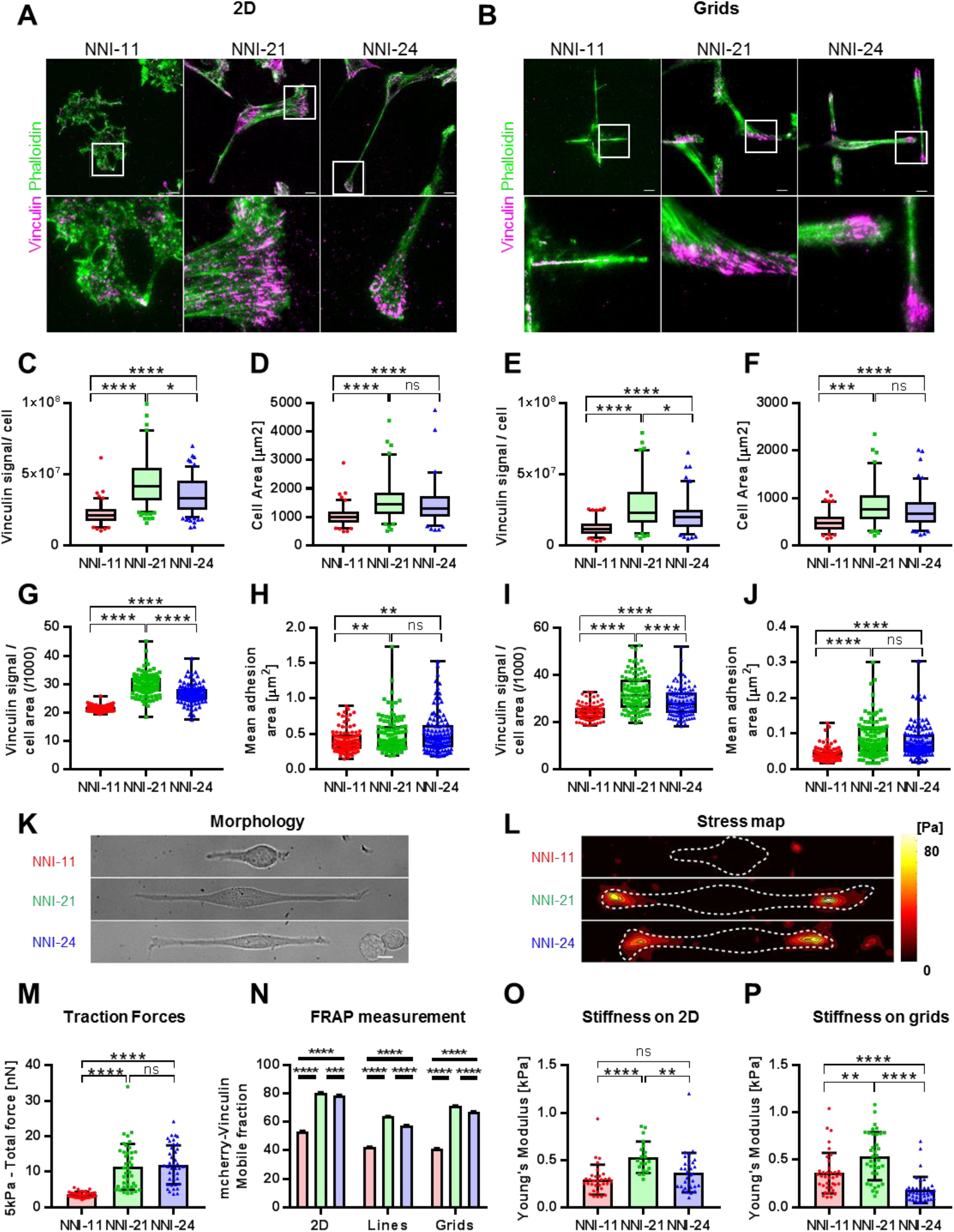
Hurdlers are stiffer and display numerous dynamic adhesions. (**A, B**) TIRF images of NNI-11, NNI-21 and NNI-24 hGPC on laminin substrates 2D and grids, fixed and stained for vinculin (magenta) and phalloidin (green). Scale bars = 10 μm. (**C-J**) Analysis of the vinculin signal on 2D (n=100, 107, 108) and grids (n=103, 111, 118). (**C**, **E**) Total Vinculin signal (A.U.) per cell (5-95 percentile). (**D, F**) Cell area in μm^2^ (5-95 percentile). (**G, I**) Total Vinculin signal reported to cell area (A.U.) (min to max). (**H, J**) Mean adhesion area in μm^2^ (min to max). (**K-L**) Morphology and stress maps of hGPC on laminin-linear substrates imprinted on 5 kPa polyacrylamide gels. (**M**) Total force (nN) of each hGPC line measured by traction force microscopy (n=40, 40, 40). (**N**) Mobile fraction (%) of mcherry vinculin measured by FRAP on 2D (n=11, 12, 12), lines (n=12, 11, 13) and grids (n=14, 15, 14). (**O-P**) Young’s modulus measured by AFM on 2D (n= 32, 20, 33) and grids (n=37, 36, 39). Number of cells (n) is given for NNI-11, NNI-21 and NNI-24, respectively. Error bars are S.E.M. for FRAP experiments and S.D. for TFM and AFM experiments.

## FMN1 expression is mechano-chemically induced in hurdlers

Cytoskeletal cohesion could be potentiated by molecules that increase the density of the cytoskeleton gel, such as formins. Formins constitute a family of 15 different proteins, containing FH1 and FH2 domains that mediate actin nucleation and polymerization (*21, 22*). Several formins have been involved in glioblastoma motility (*13, 23–25*). In accordance, we found that the formin inhibitor SMIFH2 led to a near total arrest of motility in our 3 cell lines (Fig. 3A-C, movies S5-S6). Using available antibodies, we tested the level of protein expression of seven different formins in our 3 hGPCs. We found high heterogeneity in the formin expression profile of each hGPC, that correlated with cell behavior in some cases (Fig. 3D). Strikingly, the formin 1 (FMN1) was expressed only in the hurdlers. Moreover, the expression of FMN1 increased when cells were grown on laminin substrates and increased with laminin concentration (Fig. 3E-G and fig. S5). This was the case also for FHOD3, which we found to be involved in glioma linear migration in our previous work (*13*). Since laminin has no effect on FMN1 expression when cells are not adherent (Fig. S5), the mechanical activation of one of the laminin-binding integrins such as α3, α6, or α7, is likely to be involved. Moreover, only adhesion on laminin, and not on fibronectin, can trigger this FMN1 overexpression (Fig. 3E-G). To our knowledge, this is the first observation that formin expression can be regulated by the substrate, highlighting an adaptation of the cells to their mechanical and chemical environment. Hence, the laminin located along blood vessel walls (*26, 27*) could increase the expression of FMN1 in the surrounding invading glioma cells that are escaping the tumor core. Accordingly, FMN1 expression (mRNA and protein) was higher in glioma cells located at the invasive front compared to the tumor core in xenograft and in patient tumor samples (Fig. 3H-J).

**Fig. 3.**
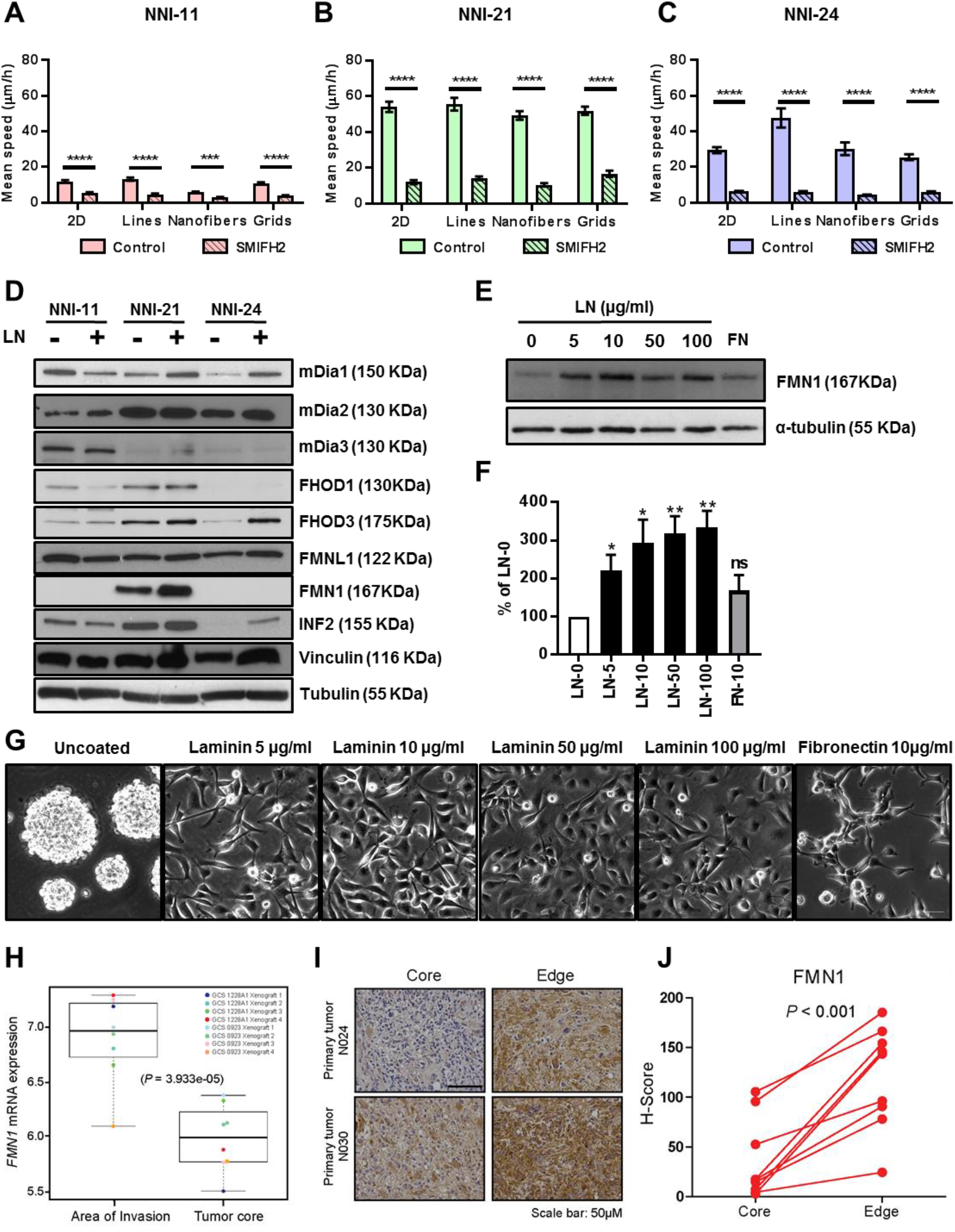
Hurdlers express FMN1. (**A-C**) Effect of SMIFH2 on hGPC migration (mean speed) on 2D (n= 40, 40, 40), lines (n= 31, 40, 41), fibers (n=18, 40, 46) and grids (n=17, 40, 56), n is given for NNI-11, NNI-21 and NNI-24, respectively. (**D**) Expression of formins in total cell extracts of the 3 hGPC cell NNI-11, 21 and 24 growing as tumorspheres without laminin coating (-) or as adherent monolayers on laminin coated plates (+). (**E**) Expression of FMN1 and tubulin in total cell extracts of NNI-21 cells growing on plates coated with laminin at concentrations ranging from 0 to 100 μg/ml or with fibronectin (10 μg/ml). (**F**) Quantification of the expression of FMN1 in each condition reported to the expression of FMN1 without laminin coating (LN0). 4 independent western blots were quantified. (**G**) Snapshots of NNI-21 seeded on plates coated with laminin or fibronectin at different concentrations. (**H**) FMN1 mRNA expression in area of glioma invasion and tumor core was evaluated in Nevo’s data set (*40*). (**I-J**) FMN1 staining were analyzed immunohistochemically in matched tumor core and edge of patient GBM tumors, representative images of two patient tumors are shown (**I**). Images were quantified using the H-score method (**J**). Scale bar is 50 μm.

## FMN1 promotes hurdling behavior

FMN1 is the founding member of the formin family (*28, 29*). It is a bona fide formin, able to nucleate actin filaments *in vitro* (*30*). FMN1 mutations caused limb deformity (*28, 29, 31, 32*). The protein was involved in cell-cell adhesion (*30*), focal adhesion formation in primary epithelial cells (*33*), dendritogenesis and synaptogenesis in hippocampal cultures (*22, 34*). All these phenotypes fitted with the hypothesis that FMN1 was a general regulator of the mechanoproperties of the cell and promoted cytoskeletal cohesion. We tested the mechanoproperties, migration and brain invasion abilities in NNI-21 after knock down of FMN1 (Fig. 4, fig. S6-S8, and movie S7). On microprinted grids, migration stopped almost completely. Knockdown cells were trapped at cross-points and adopted the shape of long crosses, suggesting that these cells had difficulty in releasing from formed adhesions to enable switching from one line to the other (Fig. 4A-C, movie S7). Adhesions appeared as thin dots spread all across the cells and did not assemble into large patches (Fig. 4D-F, fig. S6). Vinculin in these adhesions was less engaged than in control cells, as observed by FRAP measurements (Fig. 4G, fig. S8). The FMN1 knock downed cells displayed fewer traction forces and were softer than the control cells (Fig. 4H-I). Remarkably, these differences were more clearly observable on grids than on 2D substrates suggesting that the mechanical properties supported by FMN1 were necessary when cell’s cytoskeletal cohesion was challenged (Fig. S7-S8). This was further confirmed by xenograft assays in which cell invasion was challenged by the confinement of the brain environment. In this case, FMN1 - knock down cells generated smaller tumors at a lower frequency than control cells (60% of injected mice with KD cells generated tumors vs 100% in control cells)(Fig. 4J-K).

**Fig. 4.**
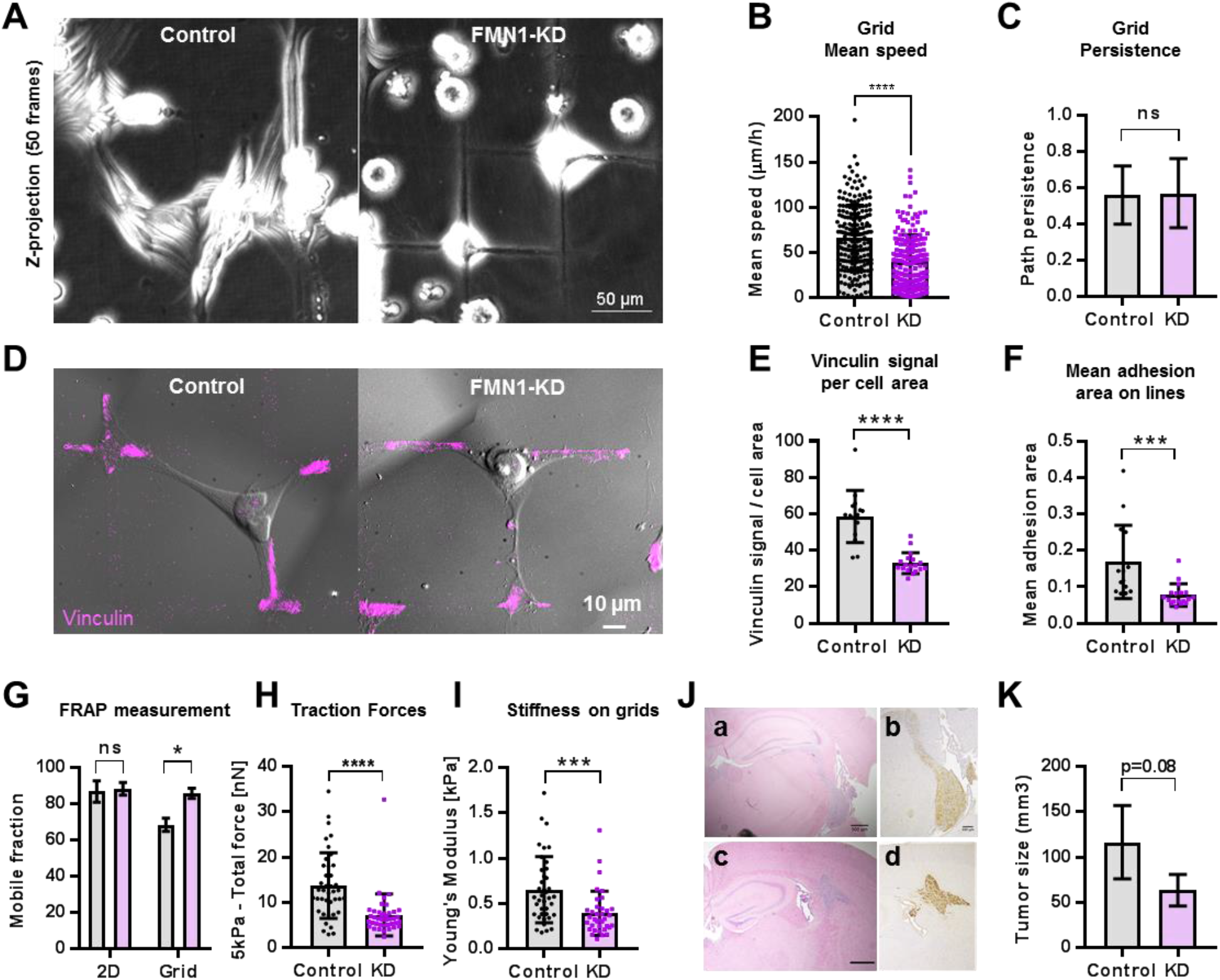
FMN1 promotes hurdling. (**A-C**) Analysis of movies of control and FMN1 knock down NNI-21 migrating on grids. (**A**) Z-projections of 50 frames corresponding to 5h of movie. (**B-C**) Average of mean speeds and path persistence (n=211, 195). (**D-G**) Analysis of adhesion in control and FMN1 knock down NNI-21, seeded on grids and lines, fixed and stained for vinculin. (**D**) Overlay of TIRF and DIC images. (**E-F**) Analysis of the vinculin signal on lines (n=15, 18). (**E**) Total Vinculin signal detected by TIRF reported to the cell area (**F**) Mean adhesion area (μm^2^). (**G**) Mobile fraction (%) of mcherry vinculin measured by FRAP on 2D (n=10, 9) and grids (n=18, 22). (**H**) Total force of control and FMN1 knock down NNI-21 measured by traction force microscopy (n=40, 40). (**I**) Young’s modulus of control and FMN1 knock down NNI-21 measured by AFM on laminin printed grids (n=39, 36). (**J**) Pictures of xenografted mouse brain slices injected with hurdlers expressing control (a,b) and FMN1 shRNAs (c,d) and sliced after 10 days. (**K**) Tumor size 10 days after injection of control or FMN1 knock down cells (n=5 mice for control and 3 mice for knock down). Number of cells (n) is given for control and FMN1-KD NNI-21, respectively.

## FMN1 organizes the cytoskeleton acting from the microtubule lattice

FMN1 could support cell cohesion by organizing the overall cytoskeleton, which consequently affects focal adhesions and cell migration. We tested this hypothesis by looking at the effects of FMN1 knock down or overexpression on the cytoskeleton and at its subcellular localization (Fig. 5, fig S9, movies 8-9). We found FMN1 in the cytosol and along the microtubules but no clear co-localization with classical F-Actin containing structures was detectable, in accordance with observation by other groups (*33, 35*). This microtubule co-localization was clear on live samples where GFP FMN1 localized along microtubules that were capped with RFP-EB3 and sensitive to Nocodazole (Fig. 5C-E, movies S8-9). FMN1 could participate in the organization of the microtubule network by separating the microtubules from each other: We found that in control cells, on gridded micropatterns, microtubules formed a spread-out network, supporting the cell architecture above non-adhesive areas, while in knock down cells microtubules assembled in tight bundles (Fig. 5A-C). FMN1 could also promote actin polymerization. When overexpressed in the non-migrating NNI-11, FMN1 led to an increase of actin filaments and vinculin-containing adhesions accompanied by a change in cell shape (Fig. 5F-G, fig. S9). The increase in actin filament density implies that FMN1 was indeed responsible for actin nucleation / elongation even though it is not co-localized with actin stress fibers. Its localization suggests that FMN1 could generate actin filaments from the cytosol and the microtubule lattice. This could increase the density of the cytoskeleton gel and enhance the cytoskeleton cohesion that helps cells to migrate and change directions readily.

**Fig. 5.**
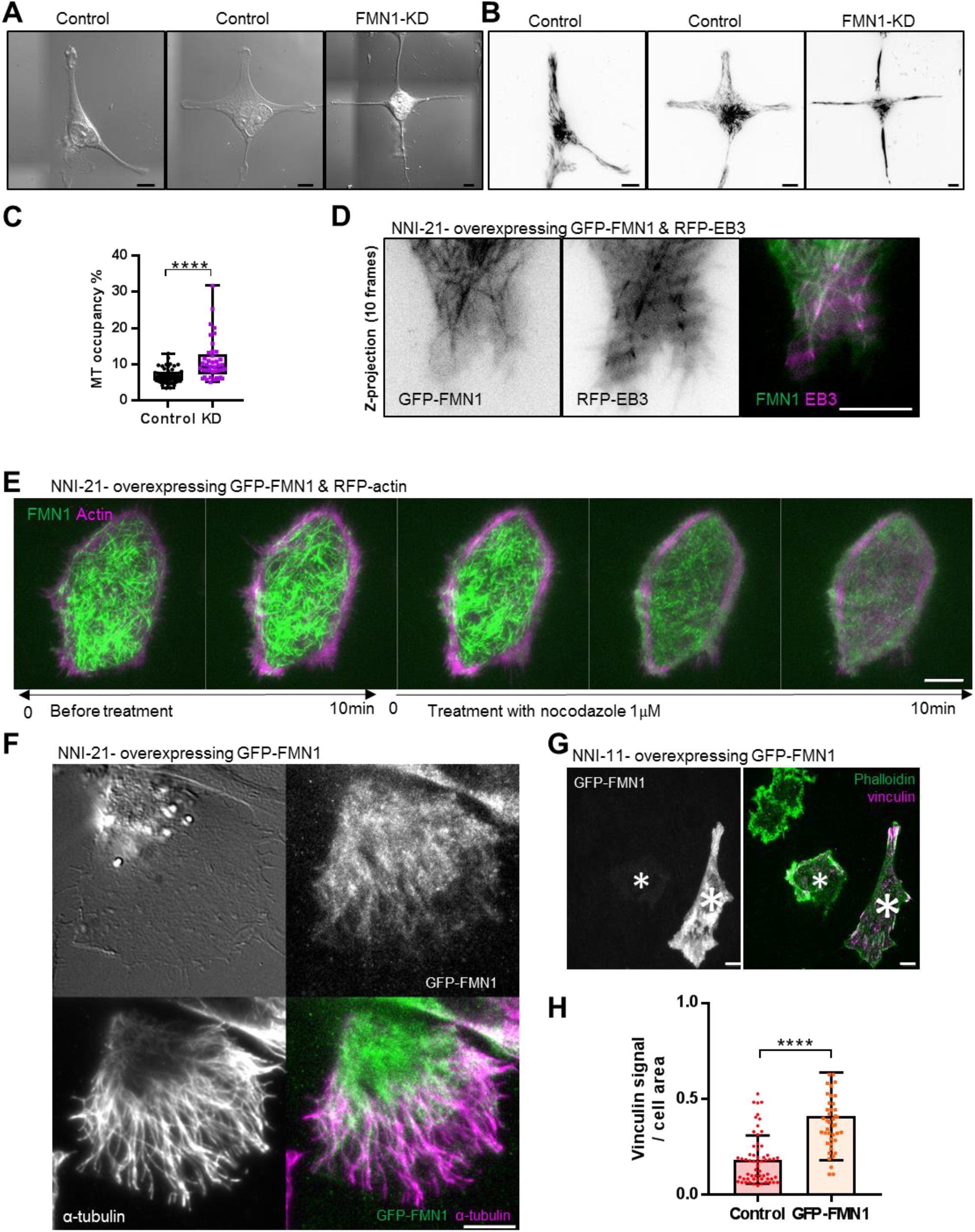
FMN1 organizes the cytoskeleton. (**A-B**) DIC (left) and TIRF images of α-tubulin (right) of control and FMN1 knock-down NNI-21 on grids. (**C**) Quantification of microtubules occupancy per cell area (n=51 and 41 cells for control and FMN1-KD, respectively). (**D**) Z projection of 10 frames corresponding to 30 sec of movie of NNI-21 hGPC expressing GFP-FMN1 and RFP-EB3. Bar is 10 μm. (**E**) Montage (10 min before treatment, and 10 min after) showing the effect of Nocodazole on the GFP-FMN1 tubules (Frequency: 5 min) extracted from TIRF movie of hurdlers expressing GFP-FMN1 and mCherry-Actin. (**F**) TIRF images of hurdlers expressing GFP-FMN1 fixed and stained for tubulin (magenta). (**G**) TIRF images of NNI-11 hGPC expressing GFP-FMN1 fixed and stained for vinculin (magenta) and phalloidin (green). Scale bars = 10 μm. (**H**) Quantification of the total vinculin signal per cell (n=60 and 43 cells for control and GFP-FMN1 expressing cells, respectively)

## Conclusions

We have reported direct measurements of several biophysical parameters of hGPCs. hGPCs display low traction force (ranging from 5 to 20 nN) and a low elastic modulus (0.2-1 kPa), in accordance with previous work on glioma stem cells, neurons and brain tissues (*2, 36–38*). We find that three different hGPCs have markedly different biophysical parameters and yet all cause severe disease states. Tumor progression is very different for the different cells and is consistent with their biophysical properties. In the case of NNI-11, the cells migrate slowly, generate low traction forces and are soft, which is consistent with their low invasive potential. In contrast, NNI-21 cells have the most rapid and random motility with the largest traction forces and most rigid cytoplasm, consistent with their rapid invasion. The NNI-24 cells are intermediate in rigidity and traction forces, but move in a highly directional fashion at a slower rate. The NNI-21 and NNI-24 cells have different modes of motility, hurdling and gliding, respectively, that involve specific mechanical parameters and specific molecular players. Differences between hurdlers and gliders are readily detected on gridded micropatterns by the temporal stack projections of the movies, without the need for individual cell tracking. Thus, the type of glioblastoma can be readily determined from the biophysical characterization. This type of design could be used in clinical settings to rapidly screen for motility modes of tumor cells and detect highly dynamic cells. We suspect that other types of motility will emerge when more hGPCs will be analyzed. Hurdling appears as a highly aggressive mode, in which cells can rapidly explore a vast substrate area and change direction without slowing down. In the brain environment, hurdlers could swap from one brain vessel to another and quickly invade the host vasculature. Hurdlers present an improved cytoskeletal cohesion (reflected by a higher stiffness), which allows cells to form highly dynamic adhesions and increase their adhesive strength and their migration capabilities (*39*). The high level of cell mechanics regulators, such as FMN1on laminin substrate as well as at the invasive front in patient tumors, can provide the cells with a robust cytoskeletal cohesion by organizing the microtubules and promoting actin polymerization. These adaptive mechanoproperties improve the overall cellular fitness that is advantageous for glioma navigation in the brain.

## Acknowledgments

We are grateful to the IFOM imaging facility personnel, in particular D. Parazzoli and M. Garrè for technical support, the IFOM histopathology facility personnel, in particular F. Pisati, the IFOM cell culture facility personnel, the MBI microfabrication facility personnel, in particular Sree Vaishnavi Sundararajan. We thank Marco Foiani and Giorgio Scita (IFOM), Virgile Viasnoff (MBI), Marc-Antoine Fardin (Institut Jacques Monod), Nir Gov (Weizmann Institute of Science, Israel), Tania Dini and all the members of Gauthier’s, Scita’s and Maiuri’s groups for helpful discussions and critical comments on the manuscript.

## Authors contributions

N.C.G. and P.M. conceptualized the research project. M.C. and P.M. developed the computer codes and algorithms. P.M., M.C., Y.K.C., K.H., A.G., Q.L., C.R., P.M., C.T. and N.C.G. analyzed the data. P.M., M.C., Y.K.C., K.H., A.G., Q.L., C.R., B.T.A.and N.C.G. performed the experiments. M.P.S., M.B., C.T., G.P., and N.C.G. provided the resources. PM wrote the original draft. All authors reviewed and edited the manuscript. M.B., M.P.S., G.P., C.T., N.C.G., supervised the research activity.

## Data and materials availability

all data is available in the manuscript or the supplementary materials.

## List of Supplementary Materials

Materials and Methods

Fig S1-S9

References (41-44)

Movies S1-S9

