## Supplemental material for "Adaptive mechanoproperties characterize glioblastoma fitness for invasion"

### Supplementary Materials

### Materials and Methods

#### *Stereotaxic Intracranial Implantations of NOD/SCID Mice*

Animal experimentation was performed according to protocols approved by the Institutional Animal Care and Use Committee of the National Neuroscience Institute of Singapore. Implantation was carried out as previously described (*41*), using NOD/SCID gamma mice (NOD.Cg-Prkdcscid Il2rgtm1Wjl/SzJ, The Jackson Laboratory). The following coordinates were used: antero-posterior=+1.0 mm; medio-lateral=+2.0 mm; dorso-ventral=-2.5 mm. Mice were euthanized by means of transcardiac perfusion with 4% paraformaldehyde upon presentation of neurological deficits with ataxia, cachexia, lethargy, or seizure. Hematoxylin and eosin staining was performed on 5-µm-thick paraffin sections.

#### *Immunohistochemistry*

Tissue sections were stained with anti-FMN1 antibody (1:200, Sigma, # HPA005465). For quantitative analysis, the percentage of stained tumor cells and intensity of staining were evaluated under high-power fields (400x) on tissue sections using optical microscopy. Immunopositivity for the FMN1 protein expression was assessed by means of a semiquantitative scoring method (H-score)(*42*). H-scores were then derived from both the staining intensity (scale of 0-3) and the percentage of positive cells (0-100%), which, when multiplied, generated a score ranging from 0 to 300. Briefly, the percentage of weakly stained cells was multiplied by one plus moderately stained cells multiplied by two plus strongly stained cells multiplied by three. At least 5 random fields were counted, and scoring was performed blinded to clinical data.

#### *Cell culture and transfections*

Graded brain tumor specimens were obtained with informed consent, as part of a study protocol approved by the SingHealth Centralised Institutional Review Board A. Human glioma propagating cells (hGPCs) were isolated and cultured, as described previously (*41*), as tumor spheres in high glucose DMEM/F12 (1:1) supplemented with sodium pyruvate, non-essential amino acid, penicillin/streptomycin, glutamine, B27 supplement (Invitrogen), bFGF (20 ng/ml), EGF (20 ng/ml) (PeproTech), and heparin (5 µg/ml) (Sigma). For transfection and migration assays, hGPCs were cultured as monolayers on laminin (10 µg/ml) coated petri dishes for 3-5 days before transfection. HGPCs transfections were performed with a Neon electroporator (Invitrogen) as per manufacturer's recommendations. For *in vivo* experiment, NNI-21 transfected with control and FMN1 shRNA, single transfected clones were selected in geneticin (Invitrogen).

#### *Micropatterning*

Laminin solution at 50 µg/ml diluted in dPBS was used for micro-contact printing. 7 µm lines and grids were printed on non-culture-treated petri dishes using microcontact printing, and passivation was done with pluronic F127 (0.2% solution) as previously described (*13*). To image F-actin, FMN1 and vinculin, stamps were performed on silanized glass-bottom dishes and passivation was done with PEG-PLL (0.1% solution in HEPES 10mM). HGPC grown on laminin-coated plates were detached with trypsin, counted, seeded on patterns and incubated at 37 C. Cell imaging typically started within the following hour.

#### *Microscopy*

Phase contrast, Differential Interference Contrast (DIC), Epi-Fluorescence and Total Internal Reflection Fluorescence (TIRF) microscopy of fixed and live specimens were performed on a Leica AM TIRF MC system equipped with temperature, humidity, and CO2 control. Long term imaging was done using a 10X objective (Leica HCX PL FLUOTAR 10x/0.30NA PH1 Objective). Acquisitions were typically obtained over a period varying from 12 to 24 h (1 image/6min). For TIRFM, two different objectives were used: HCX PL APO 63X/1.47NA oil immersion and HCX PL APO 100X/1.47NA oil immersion. The lasers used for fluorochromes excitation were 488nm, 561nm, 635nm with specific dichroic and emission filters for each wavelength. The microscope was controlled by Leica Application Suite AF software (Ver. 2.6.1.7314), and images were acquired with an Andor iXon DU-8285_VP camera.

#### *Quantification, tracking and statistical analysis*

Cells bodies were manually tracked using the manual tracking plugin of Fiji (ImageJ), coordinates of the tracks were used to calculate the mean speeds (µm/h) and persistence as described in Maiuri (*43*).

Vinculin quantification of the fixed samples was done using a custom-written code in Fiji (Image J) as followed: 1) the cell edge, which was used to calculate the cell area, was identified from the DIC channel. 2) all the adhesion signal from the vinculin channel was isolated by clearing all the signal outside the cell edge. 3) Focal adhesions were isolated by either sharpening or smoothing the image followed by thresholding using the Shanbhag method built-in in Fiji (ImageJ). 4) The binarized images obtained were used to calculate the mean adhesion area, mean adhesion circularity, and adhesion number for every cell. 5) Mean vinculin signal was calculated considering its raw integrated density normalized on the cell area.

Vinculin quantification of the live samples was done using a custom-written macro in Image J as followed: to quantitatively characterize the half-life of focal adhesions, cells were segmented, and vinculin signal was binarized. The binarized movie was then de-speckled, and the first and last 10 frames were removed to avoid bias. The obtained de-speckled shortened movie was saved as an image sequence and used for the following analysis. 2D image correlation was used to compute the normalized cross-correlation (NCC) coefficients between the first frame and all the remaining frames. A fast turnover in focal adhesions returns low values of NCC (lower similarity in image patterns), and slow turnovers return high values of NCC (higher similarity in image patterns). The built-in MATLAB (Mathworks) function for 2D NCC was applied as follows:

C = normxcorr2(FirstFrame, NthFrame)

N – 1 matrixes were thus obtained, being the first matrix the NCC coefficients between the first and the second frame (NCC1-2), the second matrix the NCC coefficients between the first and the third frames (NCC1-3), etc., where N is the total number of frames. The maximum value of each matrix was then extrapolated and normalized to the maximum of NCC1-2. Their trend overtime was fitted to a one-phase decay model in Prism 8 (GraphPad) to extrapolate the half-life of focal adhesions.

Protein level quantification from western blot experiments was done with the gel analysis tool from Image J.

Prism 8 (GraphPad) was used for statistical analysis. P-values were calculated using unpaired t-tests. Stars above graph bars indicate P values: (* indicates $P<0.1$, ** $P<0.01$, ***$P<0.001$ and **** $P<0.0001$).

#### *Immunofluorescences*

For vinculin and phalloidin staining, cells grown on glass-bottom dishes (coated with laminin or micropatterned) were fixed for 15 min in 4% paraformaldehyde in PBS and then neutralized using 10 mM $NH_4Cl$ in PBS for 15 min. For microtubule staining, cells grown on glass-bottom dishes (coated with laminin or micropatterned) were fixed for 5 min in methanol at -20°C. After fixation, cells were washed three times with PBS, blocked and permeabilized using PBS containing 0.1% Triton X-100 and 0.5% BSA for 15 min. Cells were typically incubated for 1h at room temperature in primary antibodies diluted in blocking buffer, washed 3 times with PBS (5 min each time), incubated for 45min at room temperature in secondary antibodies and phalloidin, diluted in blocking buffer, washed 3 times with PBS (5 min each time), and kept at 4°C in TBS containing 0.1% sodium azide until imaged.

#### *Soft micropatterning*

Laminin line patterned PAA hydrogels were microfabricated as described in Henning (*20*). Briefly, 32-mm glass coverslips (VWR) were plasma-treated for 30 s and incubated for 30 min at room temperature (RT) with poly-l-lysine–grafted–polyethylene glycol (0.1 mg/ml, pLL-PEG, SuSoS) diluted in Hepes (10 mM (pH 7.4), Sigma-Aldrich). After washing in dPBS (Life Technologies), the pLL-PEG–covered coverslip was placed with the polymer brush facing down onto the chrome side of a quartz photomask (Toppan) for photolithography treatment (5-min ultraviolet-light exposure, UVO Cleaner Jelight).

Subsequently, the coverslip was removed from the mask and coated with laminin (50 µg/ml) (Invitrogen), and Hylitefluor $488^{TM}$ labeled laminin (10 µg/ml) (cytoskeleton, #LMN01) diluted in dPBS for 30 min at RT. In the meantime, a premix of acrylamide (Sigma-Aldrich), N,N-methylenebis (acrylamide) (Sigma-Aldrich) and dPBS was prepared with a ratio for a final Young modulus of 5 kPa and degassed for 20 min. Fluorescent nanobeads (dark red, F-8807 PS Invitrogen) were added to the premix, and the dispersion was sonicated for 5 min (Bandelin Sonorex). To initiate polymerization, 1 µl of ammonium persulfate and 1 µl of tetramethylethylenediamine were added to 165 µl of premix and vortexed. A drop of 47 µl was immediately placed onto the protein-coated glass coverslip and covered with a previously silanized glass coverslip. Silanization was done beforehand with 100% ethanol containing 0.0035% (v/v) PlusOne Bind-Silane (GE Healthcare Life Science) and 0.0035% (v/v) acetic acid (Sigma-Aldrich). After 30 min of polymerization at RT, the sandwiched coverslips were emerged in double-distilled water (ddH2O) and separated with a scalpel. The PAA hydrogel patterned with laminin attached to the silanized coverslip was rinsed in dPBS and mounted in imaging chambers. Cells were seeded and imaged the following day for traction force measurements.

***Traction force measurements***

Force measurements were conducted using an inverted microscope (Nikon Ti-E) with a Zyla sCMOS camera (Andor) and a temperature control system set at 37°C as described in Henning (*20*). Images of fluorescent beads within the stressed and relaxed polyacrylamide substrate were taken before and after detachment of the adherent cell, respectively. The displacement field analysis was done using a homemade algorithm based on the combination of particle image velocimetry and single-particle tracking (*20*).

***Fluorescence Recovery After Photo-bleaching (FRAP)***

hGPC expressing mCherry-Vinculin were imaged 48 hours after transfection with a 100× objective and analyzed with the FRAP module of an Olympus Spinning Disk Unit equipped with a FRAP module. Circular regions of interest of 2 µm diameter were photo-bleached with the 405 nm laser at 100% intensity, and post-bleach images were followed with 15 to 20% laser intensity for 100 frames (1 frame every 0.6 seconds). FRAP data were analyzed by the Fiji plugin “FRAP Profiler” and curves fitted to the one-phase association equation by the software Graphpad Prism without pre-determined constraints:

a) I= I0+Imax*[1-e-(k)*(t)]

Where I is the relative intensity compared to the pre-bleach value, k1 represents the association rate, and t the half time recovery expressed in seconds.

***Atomic force microscopy***

AFM indentation was carried out using JPK NanoWizard3 mounted on an Olympus inverted microscope. The protocol was adapted from a previous study (*44*). A modified AFM tip (NovaScan, USA) attached with 10 µm diameter bead was used to indent the center of the cell. The spring constant of the AFM tip cantilever was ~0.01 N/m. AFM indentation loading rate is 0.5 Hz with a ramp size of 3 µm. AFM indentation force was set at a threshold of 2 nN. The data points below 0.5 µm indentation depth were used to calculate Young’s modulus to ensure small deformation and minimize substrate contributions (*44*). The Hertz model is shown below:

$$F = \frac{4}{3}\frac{E}{(1-v^2)}\sqrt{R\delta^3} \qquad (1)$$

where F is the indentation force, E is the Young’s modulus to be determined, ν is the Poisson’s ratio, R is the radius of the spherical bead, and δ is the indentation depth. The cell was assumed incompressible, and a Poisson’s ratio of 0.5 was used.

***Reagents***

Laminin (23017015) was obtained from Invitrogen and kept at -80°C. After thawing at room temperature, laminin was aliquoted and kept at 4°C. Fibronectin (11080938001) was from Sigma-Aldrich. Formin inhibitor SMIFH2 was from Sigma-Aldrich (S4826-5MG). After resuspension, in DMSO (Stock concentration 10mM), aliquots were kept at -80°C for single use. Nocodazole (M1404), Pluronic® F-127 (P2443-250G), Hexamethyldisilazane (440191), were from Sigma-Aldrich. Alexa Fluor 568–phalloidin was obtained from Invitrogen. Sylgard 184 silicone elastomer kit (1064291) was from Dow Corning. PLL-PEG (PLL(20)-g[3.5]- PEG(2) was from SuSoS.

Mouse anti vinculin clone hVIN-1 (V 9131), rabbit anti-FHOD1 (SAB4200147), rabbit anti-FHOD3 (AV463242), mouse anti-FMN1 (SAB1408502) and mouse anti-α-tubulin (T9026) antibodies were from Sigma-Aldrich. Rabbit anti-mDia2 antibodies (DP3491) and Rabbit anti-mDia3 antibodies (DP4511) were from ECM Biosciences (Versailles, KY, USA). Monoclonal anti-mDia1 antibody (610848) was from BD biosciences (Franklin Lakes, NJ, USA), rabbit anti-FMNL1 antibodies (ab97456) were from Abcam (Cambridge, England). Rabbit anti-INF2 antibodies (ABT61) were from Millipore.

Plasmids used in this study were GFP-FMN1 and myc-FMN1 (human) (obtained from Origene), vinculin-mCherry (gift from P. Kanchanawong, Mechanobiology Institute, National University of Singapore, Singapore), actin-RFP (gift from E. Lemichez, Centre Méditerrenéen de Médecine Moléculaire INSERM, Nice, France), RFP-EB3 (gift from A. Bershadsky, Mechanobiology Institute, National University of Singapore, Singapore)

***shRNA design and cloning***

shRNAs against human FMN1 were designed to target 19 nucleotide sequences and cloned into the BamH1 and HindIII cloning site of pRNAT-U6/Neo vector (GenScript corporation) which also contains a GFP marker under a cytomegalovirus promoter control. PAGE purified oligos (FMN1-human shRNA forward, 5’-GAT CCC GTA TTA CGA GAC ATC CAA ATT CAA GAG ATT TGG ATG TCT CGT AAT ACT TTT TTC CAA A-3' and reverse, 5’- AGC TTT TGG AAA AAA GTA TTA CGA GAC ATC CAA ATC TCT TGA ATT TGG ATG TCT CGT AAT ACG G -3’) were obtained from Integrated DNA Technologies. 0.5 nmol of forward and reverse oligonucleotides were diluted to 20 µl in water incubated for 4 min at 94°C and slowly cooled down to 4°C. Annealed oligos were ligated into the BamH1 and HindIII restriction sites of pRNAT-U6/Neo vector. Successful cloning was confirmed by sequencing.

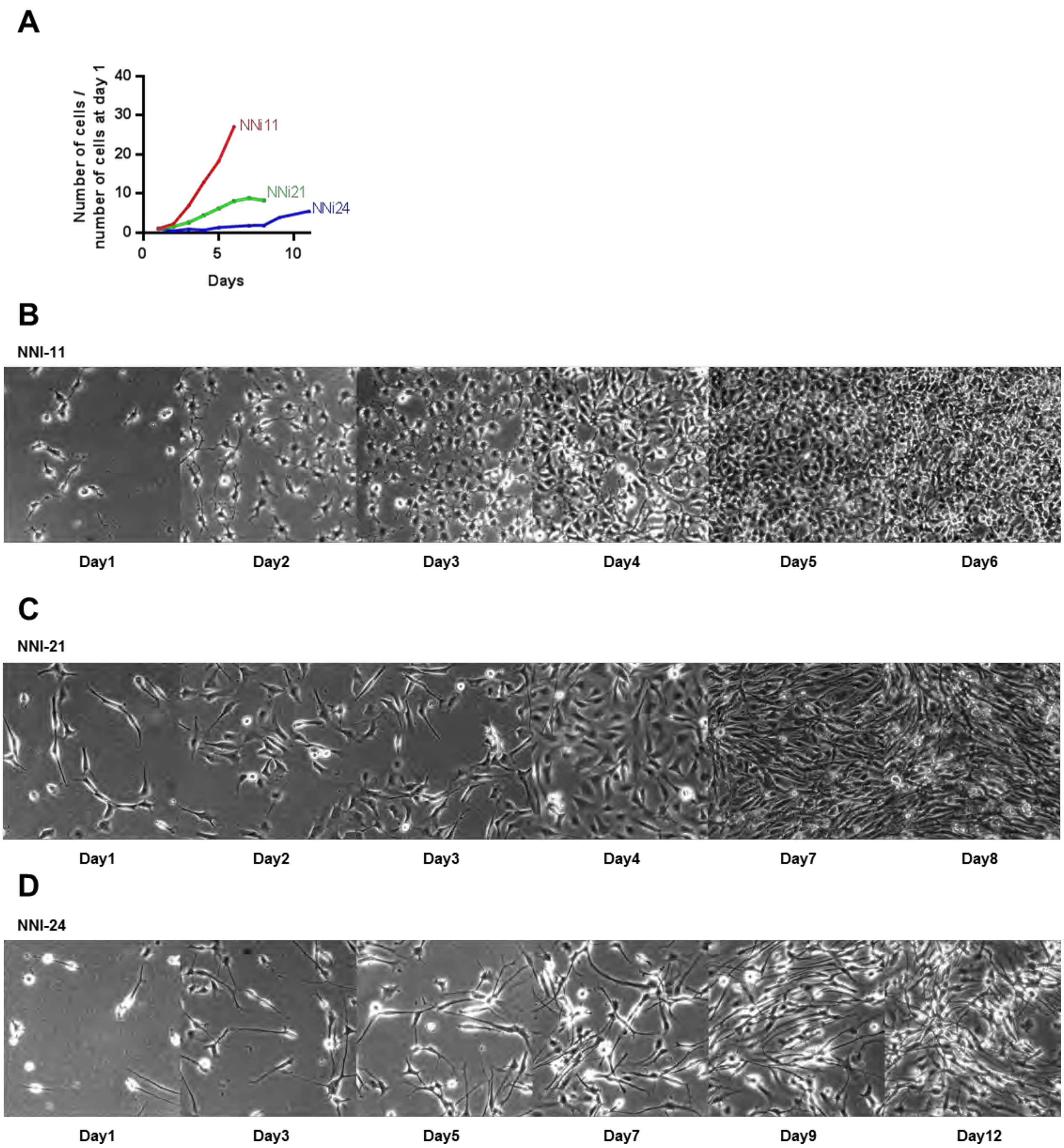


**Fig. S1**. **3 different hGPC: 3 different proliferation rates.** hGPC were cultured on 6-well plate coated with laminin. Cells were trypsinized and counted at different days in function of the cell line. The graph shows average of 2 independent experiments. (**A**)Total number of cells normalized to day 1. (**B-D**) snapshot of the cells before counting.

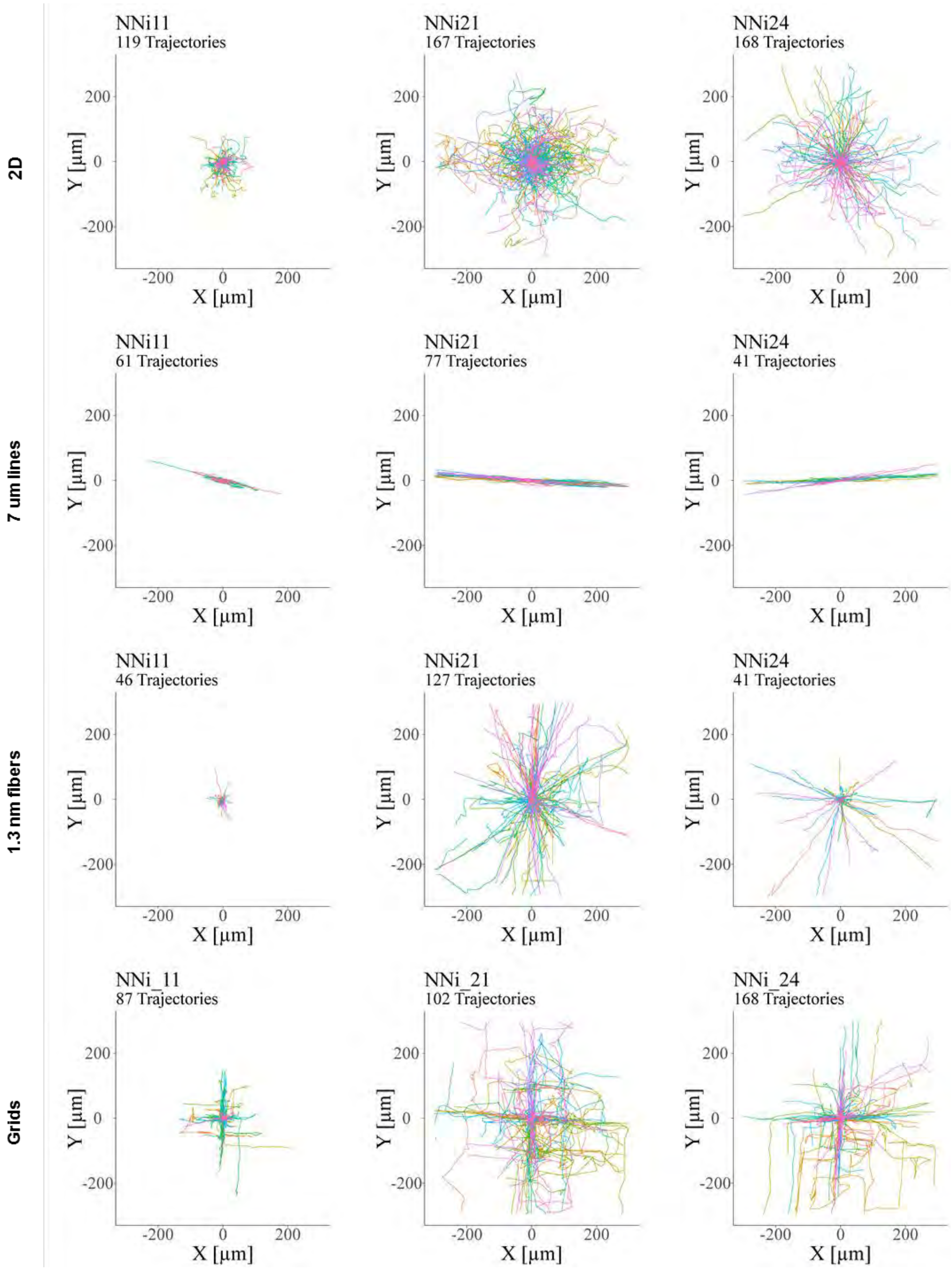


**Fig. S2**. **3 different hGPC: 3 different motility modes.** Paths traveled by the 3 hGPCs over 6 hours reported at the same origin on 2D (n= 119, 167, 168), micropatterned lines (n=61, 77, 41), nanofibers (n=46, 127, 41), grids (n=87, 102, 168). n= number of cells, is given for NNI-11, NNI-21 and NNI-24, respectively.

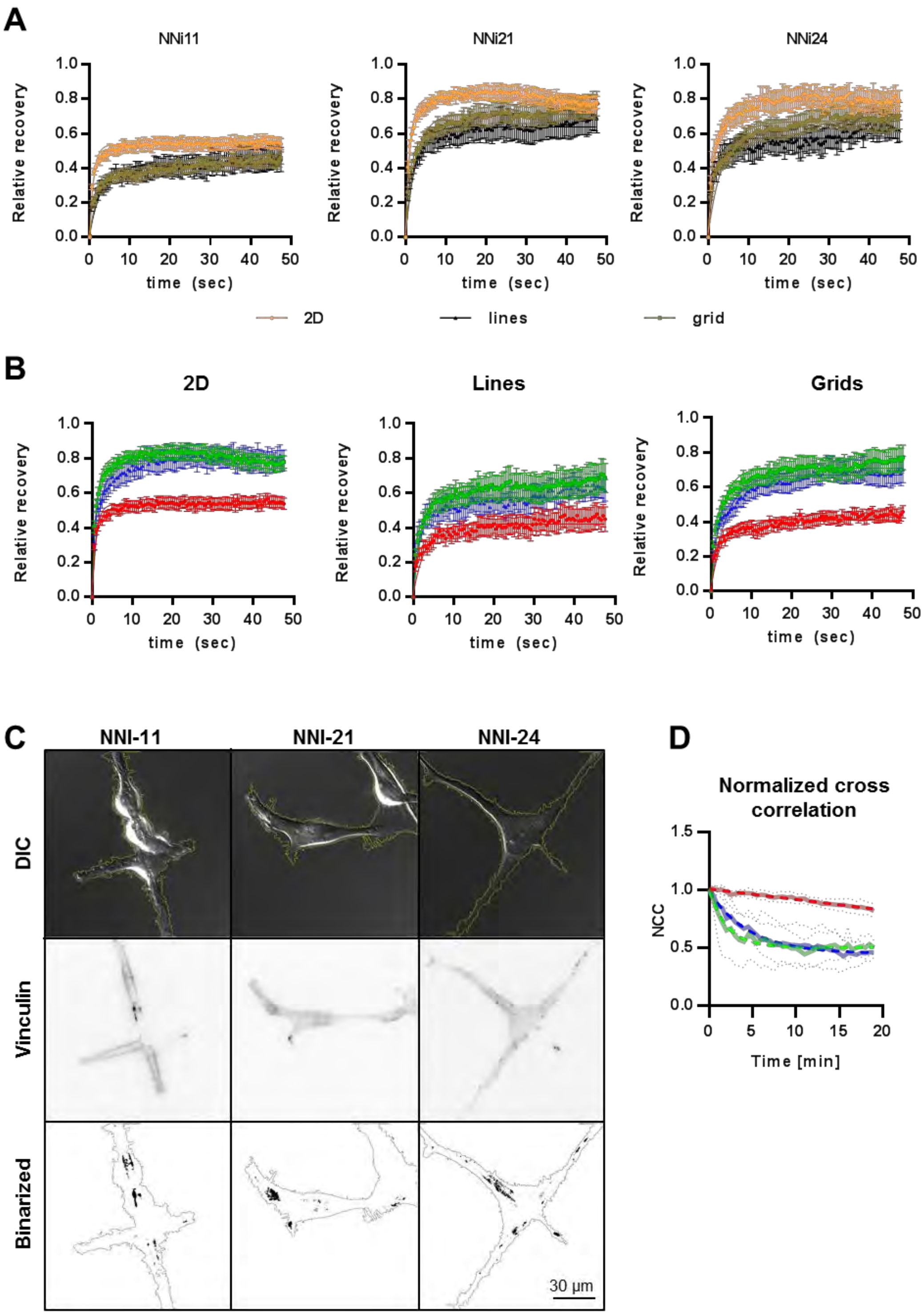


**Fig. S3**. **Comparison of the adhesion dynamics in the 3 hGPC on various substrates.** (**A-B**) Fluorescence Recovery After Photo-bleaching (FRAP) of mCherry-vinculin expressed in the 3 different hGPC on 2D (n=11, 12, 12), lines (n=12, 11, 13) and grids (n=14, 15, 14). (**A**) Comparison of the FRAP data on the 3 different substrates for the same hGPC. (**B**) Comparison of the FRAP data of the 3 hGPC for the same substrate. (**C**) DIC, TIRF and binarized images of the 3 hGPC transfected with mcherry-vinculin seeded on laminin-coated grid (snapshot extracted from Supplementary Video 4). (**D**) Normalized cross-correlation coefficients overtime for the 3 different hGPC (n=7, 5, 6). The value at each time point was calculated as max(NCC1-t) / max(NCC1-2) (see methods for details). n= number of cells, is given for NNI-11, NNI-21 and NNI-24, respectively.

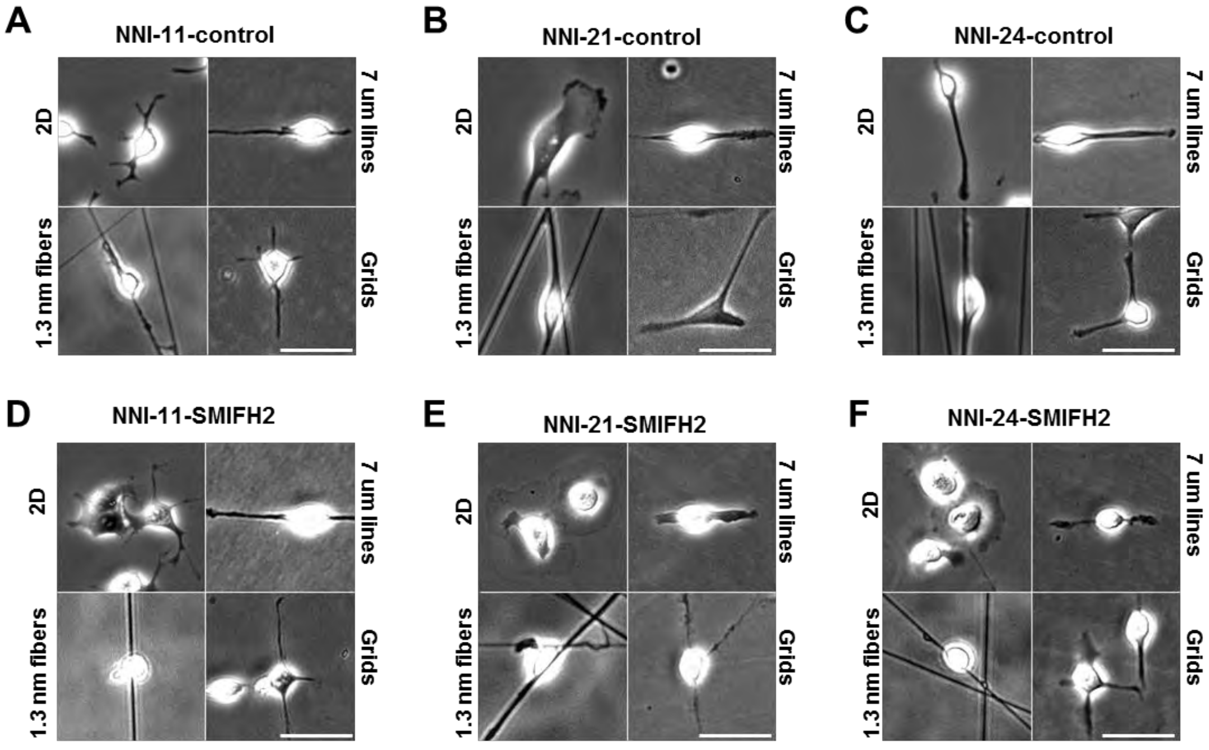


**Fig. S4**. **Effect of SMIFH2 on hGPC migration on 2D, lines, fibers and patterned grids.** NNI-11, NNI-21 and NNI-24 cells were seeded onto 4 different substrates (2D, lines, fibers and grids). After addition of SMIFH2 (final concentration 20 µM) cells were imaged in phase contrast for 6 hours (1 image / 6min). (**A-C**) Typical snapshots of NNI-11, NNI-21 and NNI-24 cells on the 4 different substrates in control condition. (**D-F**) Typical snapshots of NNI-11, NNI-21 and NNI-24 cells on the 4 different substrates 4h after the addition of SMIFH2.

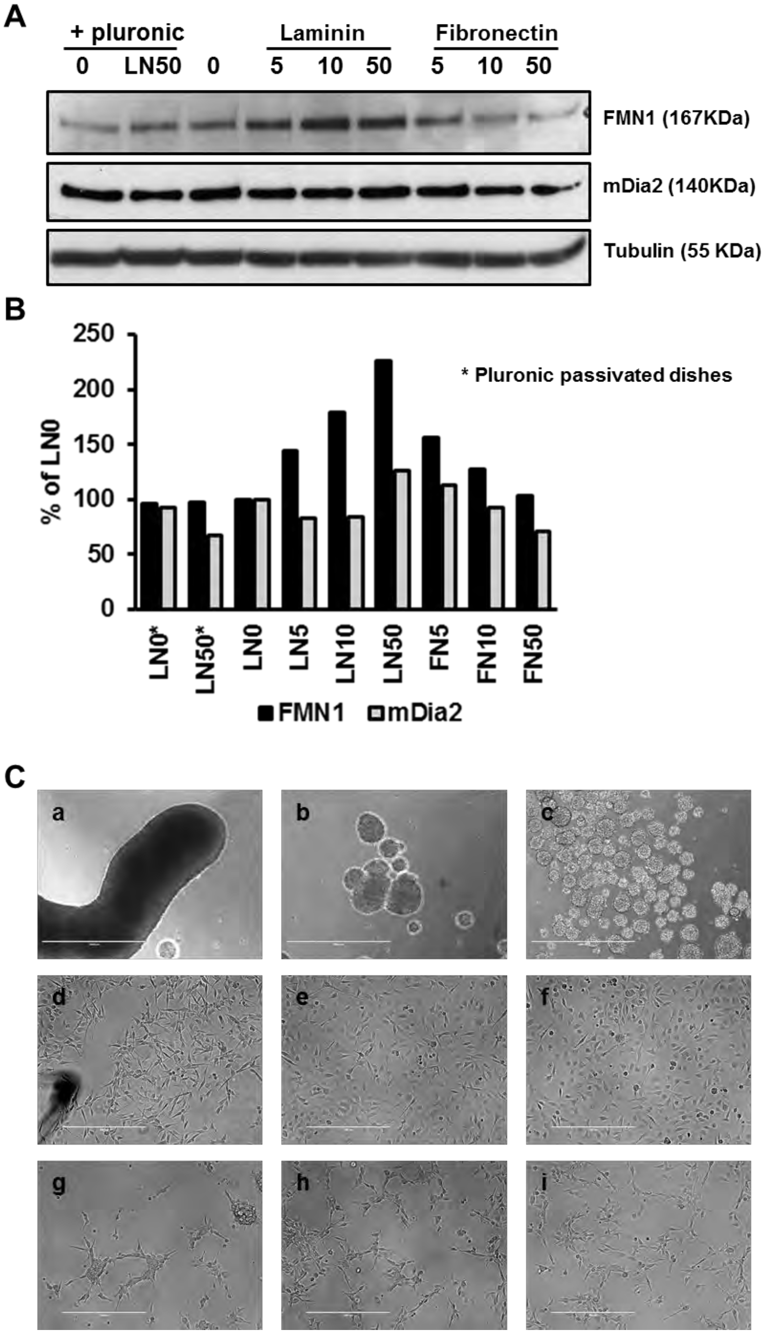


**Fig. S5**. **FMN1 expression increases on laminin substrates.** (**A**) Expression of formins FMN1 and mDia2 in total cell extracts of the hGPC NNI-21 growing on passivated dishes in the absence (LN0*) or in the presence of 50 µg/ml laminin (LN50*) and non-passivated dishes coated with laminin (LN) or fibronectin (FN) at a concentration varying from 5 to 50 µg/ml. (**B**) Quantification of the expression of FMN1 and mDia2 in each condition reported to the expression of FMN1 and mDia2 without laminin coating and no passivation (LN0). (**C**) Snapshots of NNI-21 upon the different coating conditions: **a)** passivated, no laminin; **b)** passivated, laminin 50 µg/ml; **c)** not passivated, no laminin; **d)** coated with laminin 5 µg/ml; **e)** coated with laminin 10 µg/ml; **f)** coated with laminin 50 µg/ml; **g)** coated with fibronectin 5 µg/ml; **h)** coated with fibronectin 10 µg/ml; **i)** coated with fibronectin 50 µg/ml. Bar is 1 mm.

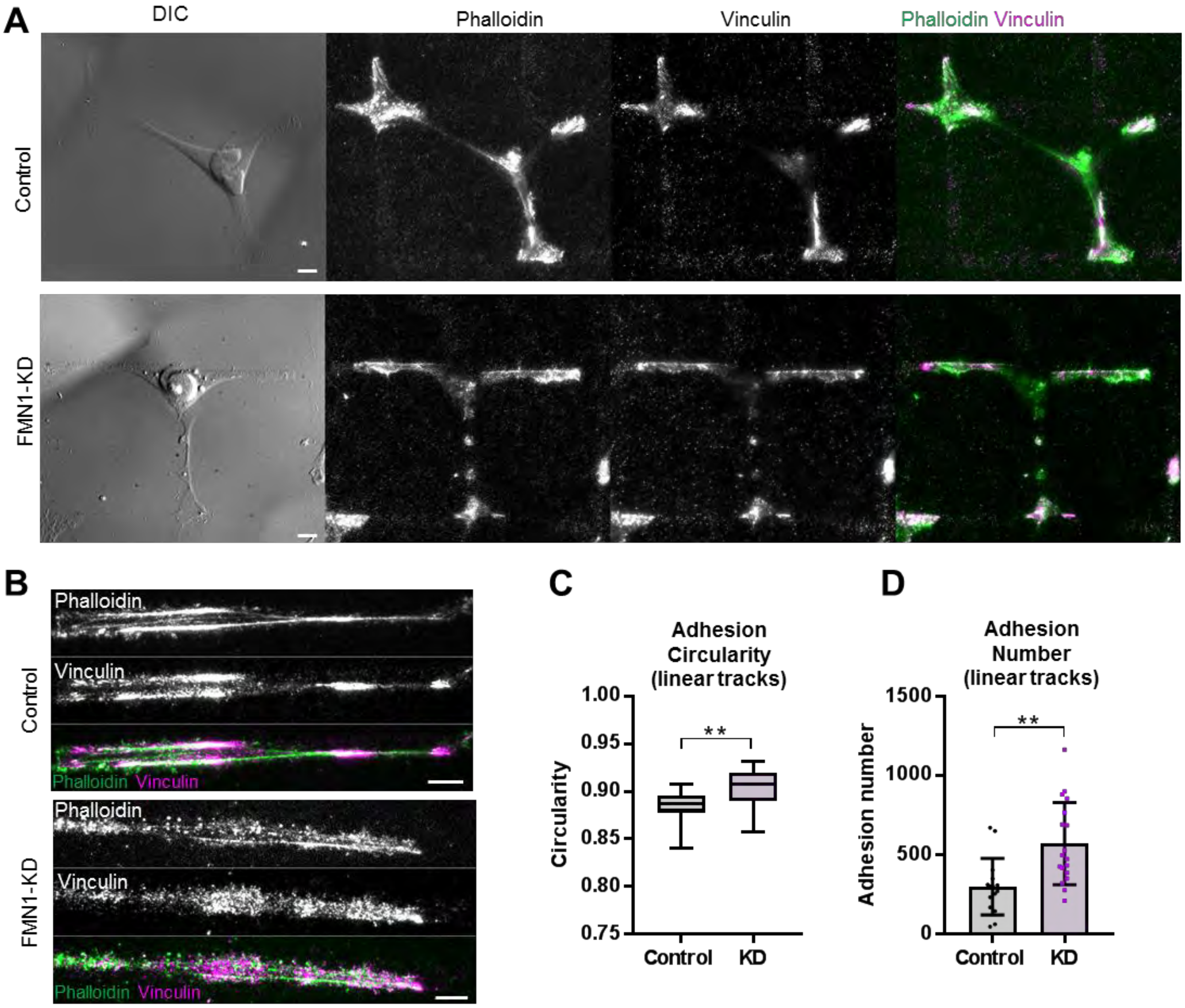


220

**Fig. S6**. **FMN1 promotes hurdling.** (**A-B**) TIRF images of control and FMN1 knock down NNI-21 fixed and stained for vinculin (magenta) and phalloidin (green), on laminin-coated grids (**A**) and lines (**B**). Scale bars = 10 µm; (**C-D**) Measurement of adhesion circularity and adhesion number on lines (analyzed from the TIRF vinculin images) (n=15 and 18 for control and FMN1 KD, respectively).

225

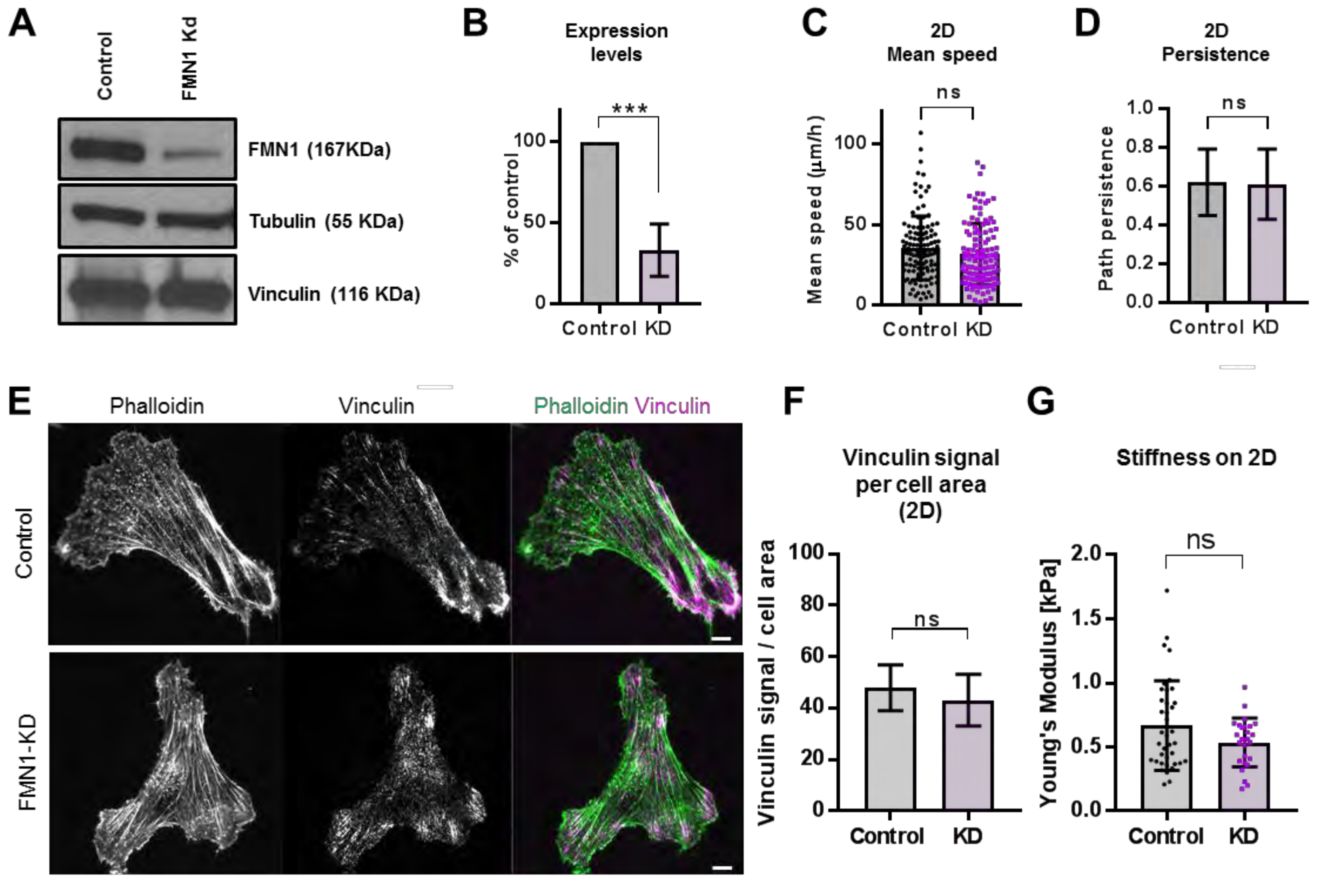


**Fig. S7**. **Effect of FMN1 knock down on 2D motility.** (**A-B**) Western blot and corresponding quantification (n=4) showing the knock down of FMN1 in NNI-21 cells; Error bars is SD. (**C-D**) Average of mean speeds and path persistence of control and FMN1 knock down NNI-21 migrating on 2D substrates (n=120 in both conditions). (**E**) TIRF images of control and FMN1 knock down NNI-21 fixed and stained for vinculin (magenta) and phalloidin (green), on 2D laminin Scale bar is 10 µm. (**F**) Total Vinculin signal detected by TIRF reported to cell area (A.U.)(n=19, 23). (**G**) Young's modulus of control and FMN1 knock down NNI-21 on 2D measured by AFM (n=38, 26). Number of cells for each measurement is indicated in table S2. Number of cells (n) is given for control and FMN1 KD, respectively.

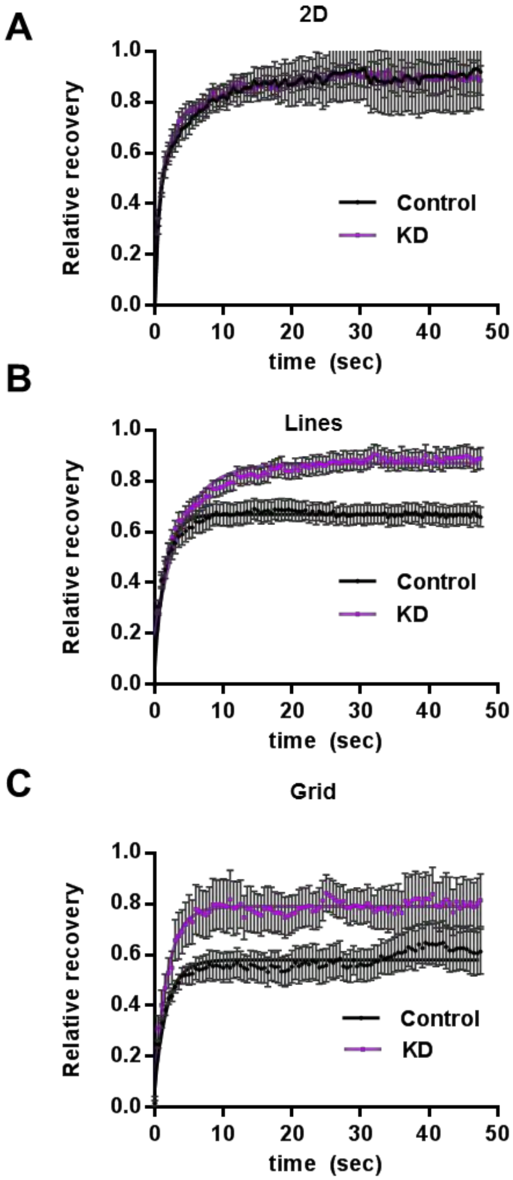


**Fig. S8**. **FMN1 stabilizes adhesions of invasive glioma on linear laminin substrates.** Fluorescence Recovery After Photo-bleaching (FRAP) of mCherry-vinculin expressed in control and FMN1 knock down NNI-21 on (**A**) 2D (n=10, 9), (**B**) lines (n=14, 11), (**C**) grids (n=18, 22). Number of cells (n) is given for control and FMN1 KD, respectively.

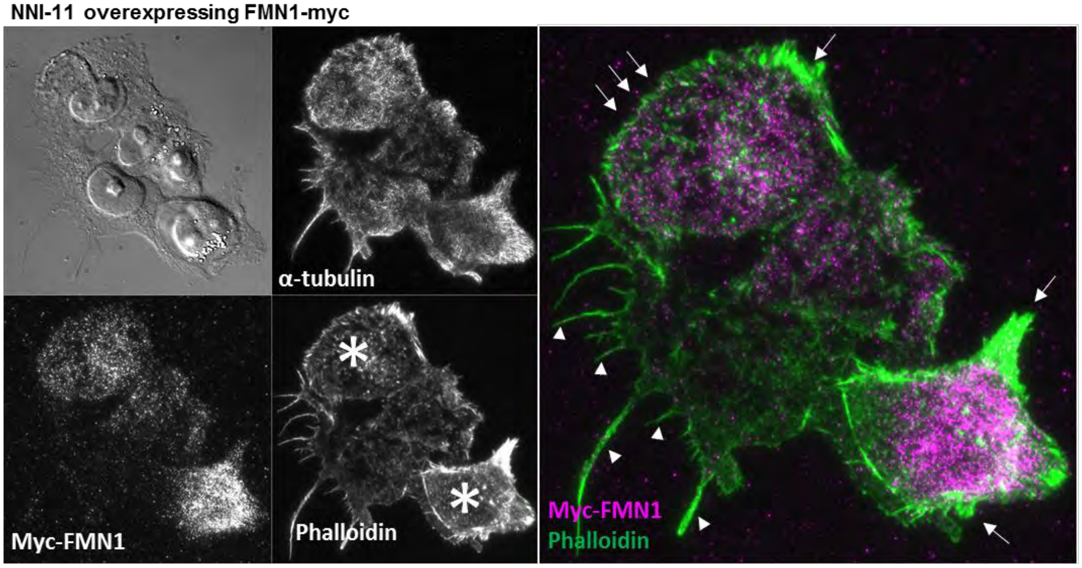


**Fig. S9**. **FMN1 promotes adhesions and stress fibers when overexpressed in NNI-11.** TIRF images of NNI-11 hGPC expressing myc-FMN1 fixed and stained for myc, vinculin (magenta) and phalloidin (green). Stars indicate transfected cells. Arrowheads indicate retraction fibers present in non-transfected cells. Arrows indicate adhesion sites and protrusions present in FMN1 expressing cells.

### Legends for Movies

**Movie S1. hGPCs migrate better on laminin than on collagen or fibronectin.**
NNI-11, NNI-21 and NNI-24 were seeded on glass-bottom multiwell plate coated with 50 µg/ml of laminin, fibronectin or collagen. 1h after plating, cells were imaged in phase contrast with a 10X objective for > 6 hours (1 image/6min). Scale bar is 50 µm.

**Movie. S2. hGPCs migrating on 2D substrates, micropatterned lines, nanofibers and micropatterned grids.**
NNI-11, NNI-21 and NNI-24 were seeded on glass-bottom dishes, suspended nanofibers, micropatterned lines and grids, all coated with laminin. 1h after plating, cells were imaged in phase contrast with a 10X objective for > 6 hours (1 image/6min). Movies are presented in order: 2D, lines, nanofibers and grids. Scale bar is 50 µm.

**Movie. S3. hGPCs migrating on micropatterned grids (large field of view).**

NNI-11, NNI-21 and NNI-24 were seeded on micropatterned grids coated with laminin. 1h after plating, cells were imaged in phase contrast with a 10X objective for > 6 hours (1 image/6min). Scale bar is 100 µm.

**Movie. S4. Adhesion dynamics of glioma cells on micropatterned grids.**
NNI-11, NNI-21 and NNI-24 expressing mcherry-vinculin were seeded on laminin grids micropatterned on glass-bottom dishes. Cells were imaged in DIC (upper panel) and TIRFM (middle panel) with a 63X objective for 2 hours (1 image/30 sec). For quantification purpose, binarized movies of the TIRF signal (lower panel) were generated. Scale bar is 30 µm.

**Movie. S5. Effect of SMIFH2 treatment on glioma cells migrating on 2D substrates, micropatterned lines, nanofibers and micropatterned grids.**
NNI-11, NNI-21 and NNI-24 were seeded on glass-bottom dishes, suspended nanofibers, micropatterned lines and grids, all coated with laminin. 1h after plating, cells were imaged in phase contrast with a 10X objective for 4 to 12 hours (1 image/6min), SMIFH2 (20 µM final concentration) was added to the dishes and cell were imaged for an extra 4 to 12h. Movies of control and SMIFH2 treated cells on the 4 settings were concatenated, starting with NNI-11 and followed by NNI-21 and NNI-24. Scale bar is 50 µm.

**Movie. S6. Effect of SMIFH2 treatment on NNI-21 cells migrating on 2D substrates.**
Zoom extracted from Supplementary Video 5.

**Movie. S7. Effect of FMN1 depletion on NNI-21 migrating on 2D substrates and micropatterned grids.**
NNI-21 cells expressing shRNA control or shRNA-FMN1 were seeded on glass-bottom dishes and micropatterned grids, all coated with laminin. 1h after plating, cells were imaged in phase contrast with a 10X objective for >6 hours (1 image/6min). Movies of control and knock down cells are shown side by side, 2D and grid conditions were concatenated. Scale bar is 50 µm.

**Movie. S8. NNI-21 expressing GFP-FMN1 and RFP-EB3 on 2D laminin substrate (TIRF imaging)**
NNI-21 cells expressing GFP-FMN1 and RFP-EB3 seeded on laminin-coated glass-bottom dishes were imaged in TIRF with a 100X objective for 10-20 minutes (1 image / 3 sec). Scale bar is 10 µm.

**Movie. S9. NNI-21 expressing GFP-FMN1 and RFP-actin on 2D laminin substrate treated with Nocodazole (1µM) (TIRF imaging)**
NNI-21 cells expressing GFP-FMN1 and RFP-actin seeded on laminin coated glass bottom dishes were imaged in TIRF with a 63X objective for 20 minutes, after ~10 minutes Nocodazole (1 µM final concentration) was added on stage and imaging continued for another 10 minutes (1 image / 10 sec).
